# Imaging traumatic brain injuries in mice with potassium channel PET tracer [^18^F]3F4AP

**DOI:** 10.1101/2025.04.23.650093

**Authors:** Karla M. Ramos-Torres, Kazue Takahashi, Kryslaine L. Radomski, Emiri T. Mandeville, Lauren Zhang, Yu-Peng Zhou, Eng H. Lo, Regina C. Armstrong, Pedro Brugarolas

**Affiliations:** Department of Radiology, Massachusetts General Hospital and Harvard Medical School; Boston, Massachusetts, USA; Department of Anatomy, Physiology and Genetics, Uniformed Services University of the Health Sciences, Bethesda, Maryland, USA

**Author notes:** **Correspondence:** Pedro Brugarolas, PhD, 55 Fruit St, Bulfinch 051, Boston, MA 02114.

**Keywords:** traumatic brain injury, demyelination, [^18^F]3F4AP, positron emission tomography, mouse study

## Abstract

**Objective:** Traumatic brain injury (TBI) can lead to secondary injury, including axon and myelin damage, which contributes to long-term neurological deficits. The PET tracer [^18^F]3F4AP, a fluorinated derivative of the FDA-approved drug 4-aminopyridine, selectively binds to voltage-gated potassium (K_V_) channels, offering a novel approach to assess TBI-related node of Ranvier disruption and demyelination. This study evaluates [^18^F]3F4AP PET in penetrating and non-penetrating TBI models.

**Methods:** Either controlled cortical impact (CCI, penetrating) or concussive (non-penetrating) TBI models were used to induce TBI in mice. Dynamic PET imaging with [^18^F]3F4AP was performed at time points of 0, 3, 7, 14, and/or 31 days post-injury (dpi), with quantitative analyses comparing tracer uptake in injured versus control regions. Luxol fast blue (LFB) staining was conducted to evaluate histological myelin loss.

**Results:** In the CCI model, [^18^F]3F4AP PET imaging demonstrated a 34% increase in tracer uptake at the injury site at 7 dpi, correlating with histological evidence of demyelination. Tracer uptake gradually declined over time, reflecting potential remyelination. The concussive TBI model showed a smaller and more diffuse increase in uptake at 7 dpi compared to CCI.

**Conclusion:** [^18^F]3F4AP PET imaging effectively detects demyelination following TBI, with very high sensitivity in penetrating injuries. These findings highlight the potential of [^18^F]3F4AP as a valuable imaging biomarker for assessing TBI progression and/or therapeutic response. Further studies are warranted to explore its clinical applicability and comparison with other imaging modalities.

## INTRODUCTION

Traumatic brain injury (TBI) occurs when an object directly impacts the brain (*i*.*e*., penetrating TBI) or when the brain moves violently within the skull due to impact, rapid deceleration, or blast waves (*i*.*e*., non-penetrating TBI). TBIs can cause transient or permanent impairments in cognitive, physical, and psychosocial functions(*1-5*), underscoring the need for effective methods to screen and monitor these injuries(*6,7*). The injury process consists of a primary phase, which involves immediate mechanical damage to blood vessels (hemorrhage) and neurons (axonal shearing), followed by a secondary phase that unfolds over months, characterized by metabolic, cellular, and molecular changes that contribute to further axon damage, cell death, tissue damage, and atrophy(*8,9*). A key component of secondary injury is myelin damage that can impair brain circuit functions(*10,11*).

Demyelination causes slowed or failed axonal conduction and renders axons vulnerable to damage. In the open skull controlled cortical impact (CCI) penetrating injury model, peak demyelination occurs around seven days post-injury(*12,13*), with progressive lesion changes over time. However, TBI with skull penetration is relatively rare in humans, and demyelination remains underexplored in the CCI model. The most common type of TBI is non-penetrating, which results in diffuse traumatic axonal injury, particularly in subcortical white matter, as demonstrated in histopathological and magnetic resonance imaging (MRI) diffusion tensor imaging studies(*14*). In preclinical models, non-penetrating TBI has been shown to cause demyelination, alterations in K^+^ channel organization, and disruption of paranodal regions flanking the nodes of Ranvier(*10,15,16*), consistent with observations in human TBI(*17*).

In the acute setting, TBI is imaged using mainly computer tomography (CT) or MRI to assess for intracranial bleeding and gross anatomical findings(*18*). However, these modalities have limited sensitivity and specificity to chronic injury-related changes that occur weeks to months post-injury. Positron emission tomography (PET) is a molecular imaging technique that can provide quantitative information about biochemical changes providing critical insights into injury progression and potential therapeutic interventions. PET has been used in rodent models of TBI to investigate changes in metabolism with [^18^F]FDG(*19-21*), neuroinflammation with the TSPO ligands [^11^C]PBR28(*22*) and [^18^F]DPA-714(*23*), changes in synaptic density with the SV2A ligand SynVestT-1(*24*), and changes in GABA_A_-benzodiazepine receptors with [^18^F]Flumazenil(*22*). Likewise, several PET tracers have been investigated in humans after mild TBI(*25*). Nevertheless, there is still a need for novel PET tracers that can image chronic changes in TBI, specially, processes that are potentially reversible like demyelination.

The PET tracer [^18^F]3F4AP, a fluorinated derivative of 4-aminopyridine (4-AP), selectively binds to K_V_ channels, which may be overexpressed and more dispersed in demyelinated axons and may provide precise visualization of chronic demyelination after TBI(*26,27*). A recent study in rats after spinal contusion injuries demonstrated a >2-fold increase in [^18^F]3F4AP binding at the injury site 7 days post injury, with sustained elevation for at least a month(*28*). Another study, detected a 40% increase in [^18^F]3F4AP binding three years after a focal intracranial TBI in a rhesus macaque, whereas other PET tracers, including [^18^F]FDG, [^11^C]PBR28, and [^11^C]PiB, failed to detect changes(*29*). These studies support the investigation of [^18^F]3F4AP as a potential biomarker to expand the detection of TBI pathology.

In the present study, we aim to investigate [^18^F]3F4AP PET imaging in two murine models of TBI. An open skull TBI model was used to further evaluate a penetrating TBI, since in a prior study [^18^F]3F4AP unexpectedly revealed a surgical brain injury in an experimental monkey(*29*). A closed skull TBI model was used for clinical relevance to the majority of cases of human mild TBI. This non-penetrating TBI model shows demyelination in the corpus callosum, changes in axonal K^+^ channels, and improvement of axon damage with acute, low dose 4-AP treatment(*15,16*). This approach will facilitate a better understanding of demyelination dynamics and help establish [^18^F]3F4AP as a translational imaging tool for TBI research and clinical applications.

## MATERIALS AND METHODS

### Compliance

All mouse procedures were approved by the Institutional Animal Care and Use Committee (IACUC) at the Massachusetts General Hospital (MGH) and the Uniformed Services University of the Health Sciences (USU). All animal studies were conducted in compliance with the ARRIVE guidelines (Animal Research: Reporting in Vivo Experiments) for reporting animal experiments.

### Penetrating TBI model

Adult female C57BL/6J mice (10-12 weeks-old, RRID:IMSR_JAX:000664) were subjected to controlled cortical impact (CCI) (n = 26) or sham surgery (n = 5) as previously described(*30*). Briefly, mice were anesthetized with isoflurane (1-2%), positioned in a stereotactic frame and a 5 mm craniectomy was performed over the right parietal cortex between bregma and lambda while keeping the dura intact. A controlled impact (impact velocity 6 m/s, depth 0.6 mm, impact duration 150 ms) was then induced with a 3 mm diameter flat-tipped impactor (Pneumatic impact controller, MGH Biomedical Engineering Facility) placed on the dural surface. Rectal temperature was maintained at 37 ºC with thermostat controlled heating pad. After the impact, the incision was sutured, and animals allowed to recover in their cages. Sham animals underwent the same procedure except for the impact. Two CCI mice and one sham were excluded due to bleeding during the procedure. The duration of the procedure was about 10 min.

### Non-penetrating TBI model

Adult male C57BL/6J mice (8-10 weeks-old, RRID:IMSR_JAX:000664) were subjected to a closed-skull TBI (n = 6) or sham (n = 6) procedures at the USU site as previously described(*16*). Briefly, after a midline incision of the scalp, a 3-mm flat tip impactor with rounded edges (Neuroscience Tools, Cat# 2520-3S) of an ImpactOne Stereotaxic Impactor (Leica Biosystems, Buffalo Grove, IL) was zeroed on the surface of the intact skull centered at bregma and the impactor set to 4 m/s with a dwell time of 100 ms and a depth of 1.5 mm from the surface of the skull. No skull fractures or bleeding were detected after the impact. After scalp closure with Gluture (World Precision Instruments, Cat# 503763), mice were placed in a warm cage to recover. Sham mice followed identical procedures but did not receive the impact. Predetermined exclusion criteria included: (a) 10% body weight loss, poor health, or abnormal behavior at any point during experimentation (n=0); (b) depressed or fractured skull after impact (n=0); (c) post-injury righting reflex (time to flip from supine position after surgery) > 2 min for shams and < 2 min for closed-skull TBI mice (n=0). Sham and TBI mice were shipped to MGH 3 days after surgical procedures following Institutional requirements for the transfer of live animals.

### Radiotracer production

[^18^F]3F4AP was produced in a GE TRACERlab Fx2N synthesizer according to previously reported procedure(*31*). Semipreparative HPLC purification was performed using a Sykam S1122 HPLC system equipped with UV and radiodetectors on a Waters C18 preparative column (XBridge BEH C18 OBD Prep Column 130 Å, 5 µm, 10 mm × 250 mm). The obtained [^18^F]3F4AP fraction (radiochemical purity > 99%, molar activity = 3.0 ± 0.9 Ci/µmol) at end of synthesis (EoS); n = 7), in 95% 20 mM sodium phosphate buffer, pH 8.0, 5% EtOH solution) passed the QC (with Thermo Scientific Dionex Ultimate 3000 UHPLC equipped with Waters XBridge BEH C18 analytical column [130 Å, 3.5 µm, 4.6 × 100 mm], 95% 10 mM sodium phosphate buffer, pH 8.0, 5% EtOH solution as mobile phase, t_R_ = 4.7 min) and was ready for injection into animals.

### [^18^F]3F4AP *PET/CT image acquisition and reconstruction*

Mice were anesthetized with isoflurane gas and intravenously administered approximately 200 µCi of [^18^F]3F4AP via a tail vein catheter. Dynamic PET data was acquired in a tri-modality PET/SPECT/CT scanner (Triumph LabPET from Trifoil imaging) or a Sedecal SuperArgus PET/CT scanner for 30 min under anesthesia (1.5 % isoflurane, O_2_ flow 2.0 L/min). PET data was reconstructed in frames (10 × 30s, 5x 60s, 4 × 300s) for generating time-activity curves (TACs). CT was acquired following PET for anatomical reference when possible. Penetrating-TBI mice were scanned at baseline (3 days before surgical procedure) and at 3, 7, 14, 31 dpi. Non-penetrating TBI mice were scanned at 7 dpi. Some mice were scanned multiple times at least 2 weeks apart.

### Data Analysis

All data analyses were performed by investigators blinded to the experimental group assignment (TBI or Sham). Data from animals with poor tracer administration (e.g., extravasation or failed tail vein catheterization) or inadequate anesthesia were excluded from the analysis.

### Standardized Image segmentation method for penetrating TBI

PET images were analyzed and visualized using AMIDE. PET and CT images were co-registered. Using CT for anatomical reference, the craniectomy site was located, and 8 mm^3^ ellipsoid regions of interest (ROIs) were drawn below the injury. An identical ROI was placed on the contralateral side of the brain for comparison. In cases where CT was not available a template CT from another mouse was used to align the brain and position the ROIs. TACs and PET average uptake values (SUVs) from 10-30 min after injection were extracted for each ROI and ratios of the signal from the ipsilateral injury ROI to that from the contralateral ROI were calculated to allow for comparison across animals.

### Individualized image segmentation method for penetrating TBI

Segmentation and quantitation for individualized ROIs was performed with Bruker’s PMOD software. PET and CT images were superimposed in the ‘Fusion’ mode on PMOD and a custom-drawn ROI was made at the site of injury based on the craniectomy site visualized on the CT. The ‘pen’ tool was then used to draw 2D contours of the ROI in adjacent horizontal slices of the brain. This process was repeated on the contralateral side of the brain as well as the cerebellum to serve as control ROIs. TACs and PET SUVs from 10-30 min after injection were extracted for the three ROI and ratios of the signal from the injury ROI to that from the contralateral and cerebellum ROIs were calculated to allow for comparison across animals.

### Individualized image segmentation method for non-penetrating TBI

Segmentation and quantitation for individualized ROIs was performed with Bruker’s PMOD software. First, the built-in Ma-Benveniste-Mirrione atlas in PMOD was adapted to include a custom injury ROI in its segmentation. This was done by identifying the coronal slice containing both the corpus callosum and anterior commissure, and delineating the corpus callosum region found to be affected by this single impact TBI in a previous study(*14*). For each dataset, the custom atlas was first transformed to fit the CT, and then the PET and CT images were superimposed in the ‘Fusion’ mode on PMOD. The cerebellum and injury VOIs were delineated, and the ratio of their activities was taken for every scan. Investigator performing the analysis was blinded to the group allocation (TBI or sham).

### Ex-vivo analysis – Luxol fast blue staining

Mice were euthanized by intraperitoneal (IP) injection of 200 mg/kg pentobarbital sodium and phenytoin sodium solution (Euthasol^®^) for histological analysis at baseline, 3, 7, 14 and 52 dpi. Brain tissue was harvested, patted dry on a paper towel and embedded in OCT in a cryostat mold and frozen on dry ice. Coronal sections (20 µm) of brain tissue were then obtained using a cryostat at -16 ºC, mounted onto microscope slides, allowed to air dry and stored at -80 ºC. For Luxol fast blue (LFB) staining, selected frozen tissue sections on microscope were stained with a standard staining protocol. Briefly, slides were incubated in 0.1% LFB in acidified 95% ethanol overnight at 56 °C. The slides were then differentiated and counterstained with 0.05% lithium carbonate and 0.1% cresyl violet solution.

### Statistical analysis

Statistical analysis of *in vivo* PET was performed using GraphPad Prism (version 10.2). Each data point in the plots represents one mouse. Descriptive statistics including mean and standard error of the mean (SEM) were calculated for each group. Two-group *t*-tests and multiple comparisons tests (*i*.*e*., ANOVA) with a significance level of α = 0.05 were used to assess differences among groups. Grouped data are mean ± SEM.

## RESULTS

### Study design

This study aimed to evaluate the potential of the PET tracer [^18^F]3F4AP to image demyelination after TBI in mice. Two previously characterized mouse models of TBI were selected to assess changes in tracer uptake in the brain after injuries of different etiologies.

The penetrating CCI model, produces cortical damage and white matter injury that includes axon and myelin loss(*13*). Female mice were used for this model to prevent post-surgical fighting that could exacerbate the wound. After establishing a timepoint for peak response (7 dpi), tracer uptake was evaluated at multiple timepoints between 0 to 31 days post injury. For each time point, a cohort of six to eight CCI mice and two sham mice was imaged, resulting in a total of 45 scans. Data from 11 out of 45 CCI-related scans were excluded due to poor tracer administration or inadequate anesthesia. The final sample sizes for analysis were: baseline (n = 5), 3 dpi (6 TBI, 1 sham), 7 dpi (6 TBI, 1 sham), 14 dpi (6 TBI, 2 sham), and 31 dpi (5 TBI, 1 sham).

Following the CCI study, [^18^F]3F4AP was evaluated at 7 dpi in a non-penetrating concussive TBI model, which is known to produce axonal damage, node of Ranvier disruption and demyelination in the corpus callosum (*15,16*). Male mice were used in this model due its prior validation in males and the absence of post-surgical fighting. This cohort consisted of six concussive TBI and six sham mice. Data from three mice were excluded due to poor tracer administration, yielding a final dataset of four TBI and five sham datasets for analysis.

Post-mortem brain tissue from both models was collected from a subset of animals at each time point and evaluated using LFB staining of myelin to validate PET imaging findings.

### [^18^F]3F4AP PET seven days after penetrating TBI

In this study, CCI in mice was used to investigate whether [^18^F]3F4AP PET can detect spared demyelinated axons post injury. Since previous reports have described peak demyelination and upregulation of K_v_ channels around 7 dpi in models of penetrating TBI(*12,13*), we first evaluated [^18^F]3F4AP at this timepoint. CCI mice underwent dynamic PET imaging for 30 minutes, followed by a CT scan for anatomical reference. PET/CT images showed an area of high focal [^18^F]3F4AP uptake at the site of injury, which corresponds to the impact location beneath the craniectomy site confirmed by the co-registered PET/CT images (**Fig 1 A**, left). This increased tracer uptake was not observed in sham animals, which received craniectomy surgery but were not subjected to cortical impact (**Fig 1A**, right). To determine whether the increase in PET signal was due to increased perfusion or increased binding, 8.3 mm^3^ ellipsoid ROIs were drawn at the epicenter of the injury below the craniectomy site and at an equidistant position on the contralateral side. TACs were extracted for the ROIs. For CCI mice, the TACs showed a similar initial peak indicating similar perfusion and a slower washout at the injury ROI compared to the contralateral side (**Fig 1B**, left). In sham mice, no apparent difference was observed between the ipsilateral and contralateral TACs (**Fig 1B**, right). Based on the TACs, we selected the standardized uptake value (SUV) from 10-30 min as a proxy of binding and quantified the signal at the injury ROI and its corresponding contralateral ROI. CCI TBI mice showed a SUV_10-30 min_ of 1.16 ± 0.16 (n = 6) in ipsi ROI and 0.86 ± 0.10 (n = 6) in the contra ROI (**Fig 1B**, left bar graph). In the sham mouse, no difference in SUV_10-30 min_ between ipsi and contra ROIs was observed (**Fig 1B**, right bar graph). When comparing the ratio in SUV of the ipsi v*s*. contralateral side, we found an SUVr _ipsi/contra_ of 1.34 ± 0.04 in CCI mice (n = 6) and 1.02 in a sham mouse indicating that the high uptake at the injury in TBI mice is due to the cortical impact and not due to the craniectomy procedure.

**Figure 1.**
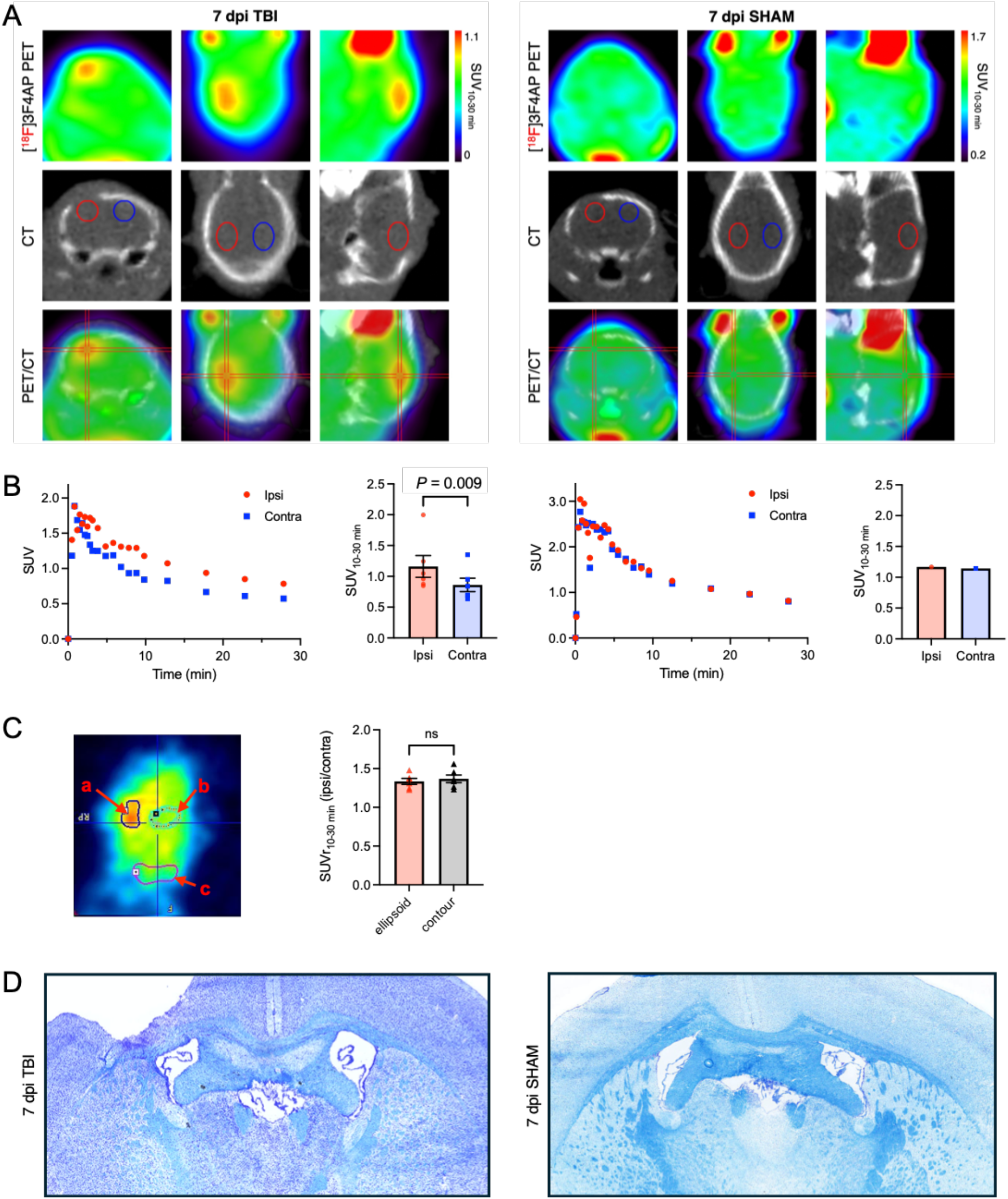
Evaluation of [^18^F]3F4AP 7 days after penetrating traumatic brain injury in mice. **A**. Representative coronal, horizontal and sagittal views of brain [^18^F]3F4AP PET/CT in a CCI TBI (left) and Sham (right) mouse. Standardized CT-guided ROIs selected are shown in CT panel in red for lesion site and in blue for contralateral region. **B**. Representative TACs extracted from the ROIs corresponding to the lesion (ipsi) and contralateral (contra) sites and corresponding quantification of standardized uptake value from 10 to 30 min (SUV_10-30 min_) for CCI TBI (*n* = 6; left panels) and sham (*n* = 1; right panels) mice. Statistical analysis was performed using a unpaired two-tailed t-test. **C**. Representative horizontal view of brain [^18^F]3F4AP PET in a CCI TBI mouse showing individualized thresholded ROIs for the lesion (a), contralateral (b) and cerebellum (c) regions and comparison of SUVr_10-30min_ using standardized (ellipsoid) *vs*. individualized (contour) ROIs in CCI mice (n = 6). Statistical analysis was performed using a paired two-tailed t-test. **D**. Selected LFB staining of a CCI TBI and sham mouse brain slices.

Due to the variable nature of the impacted area, we also analyzed the PET imaging data in a more individualized manner. To do so, an ROI for the cerebellum was drawn based on a mouse brain atlas and used as positional reference. Then, ROIs for the injured and contralateral regions were contoured based on the PET image approximately equidistant to the cerebellum ROI (**Fig 1C**, left). Using this method, the SUVr _ipsi/contra_ was 1.37 ± 0.05 for CCI mice (n = 6) and 1.06 for the sham mouse (n = 1) (**Fig 1C**, plot), which is comparable to the finding using the standardized method of ellipsoid ROIs.

Following the PET scans, brain tissue sections near the injury epicenter were collected and stained with LFB for myelin and counterstainted with cresyl violet nuclear stain. Lighter shade of LFB in the corpus callosum of CCI mice indicated demyelination and tissue damage around the impacted area (**Fig 1D**, left) which was not observed in the sham animal (**Fig 1D**, right).

### Imaging injury progression at different timepoints after penetrating TBI

After observing high [^18^F]3F4AP uptake at the injury at 7 dpi, we performed PET imaging at multiple timepoints (0, 3, 7, 14 and 31 dpi) to assess whether [^18^F]3F4AP can detect injury progression. Increased tracer uptake in the ipsilateral region was evident as early as 3 dpi, with a well-defined focal injury at 7 dpi and apparent decreases in contrast at 14 and 31 dpi (**Fig 2A**). Quantification of tracer uptake using the standardized ROI method showed a peak in SUVr _ipsi/contra_ at 1-week post injury and a return to baseline by 31 dpi (**Fig. 2B**, left plot). This suggests that there is a high concentration of demyelinated fibers in the acute (3 dpi) and subacute phases (7 and 14 dpi) and substantial remyelination and/or axonal loss by 31 dpi. In comparison, no changes in tracer uptake were observed in sham animals confirming that the higher signal is not due to the craniectomy procedure (**Fig. 3**, n = 1-2 per timepoint).

**Figure 2.**
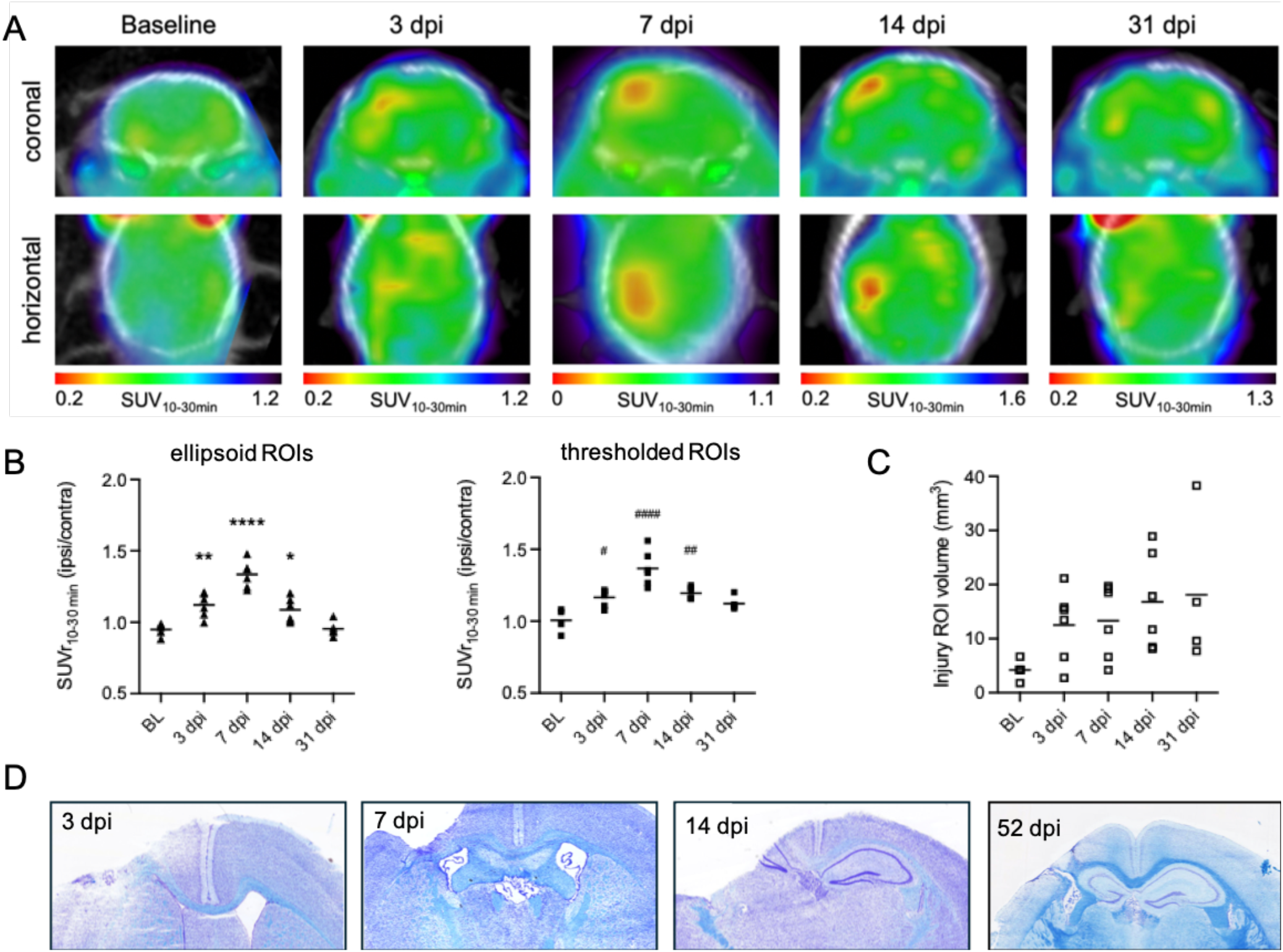
Longitudinal evaluation of [^18^F]3F4AP in the penetrating TBI model. **A**. Representative coronal and horizontal views of brain [^18^F]3F4AP PET/CT at baseline (BL), 3, 7, 14, and 31 dpi. **B**. SUVr was quantified by normalization of SUV_10-30min_ from injury ROI to contralateral ROI using standardized ellipsoid ROIs (left plot) and individualized ROIs (right plot). **C**. Comparison of volume for the individualized injury ROIs per timepoint. **D**. Selected LFB staining of brain sections from mice at 3, 7, 14 and 52 dpi showing tissue atrophy and demyelination. Statistical analysis was performed using a two-way ANOVA with Dunnett’s multiple comparison test. * # denote comparison versus BL (^**^*p* = 0.004, ^****^*p* < 0.0001, ^*^*p* = 0.02; ^#^*p* = 0.01, ^####^*p* < 0.0001, ^##^*p* = 0.003). For *in vivo* PET quantification, BL *n* = 5, 3 dpi *n* = 6, 7 dpi *n* = 6, 14 dpi *n* = 6, and 31 dpi *n* = 5.

**Figure 3.**
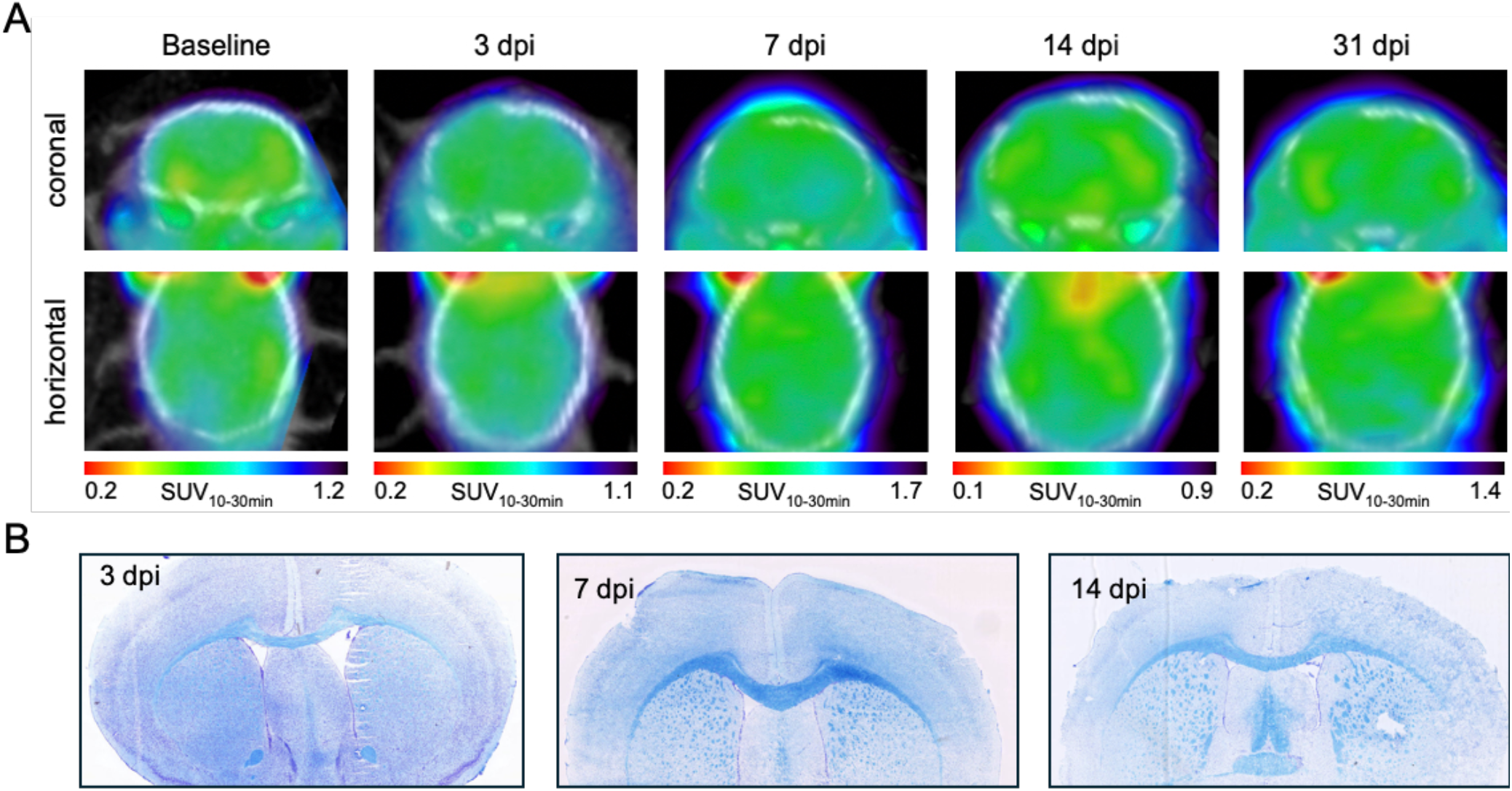
Longitudinal evaluation of [^18^F]3F4AP in open head sham mice. **A**. Representative coronal and horizontal views of brain [^18^F]3F4AP PET/CT at baseline, 3, 7, 14, and 31 dpi. **B**. Selected LFB myelin staining of brain sections from animals at 3, 7 and 14 dpi showing no tissue atrophy and demyelination.

Given the dynamic and heterogenous nature of the injury, we also applied the individualized (thresholded) ROI method to account for changes in lesion size and shape over time. Using this method, SUVr _ipsi/contra_ showed very similar results as using standardized ROIs (**Fig. 2B**, right plot). Even though the contrast at the injury was reduced over time, no significant changes were observed in the volume of the ROI that would suggest injury spreading (**Fig. 2C**).

Finally, LFB staining of brain slices taken from representative animals euthanized at 3, 7, 14 and 52 dpi revealed substantial tissue atrophy and demyelination at 3, 7 and 14 dpi and apparent remyelination by 52 dpi, consistent with the changes observed by PET and previous LFB staining results reported for this model(*12*).

### [^18^F]3F4AP PET seven days after non-penetrating TBI

The observed [^18^F]3F4AP detection of a focal injury after a penetrating TBI was next compared with signal in a model of non-penetrating TBI. We chose to compare these two models at 7 dpi as this time point showed the highest signal after penetrating injury (**Fig. 2**). To do so, we used a previously characterized closed-skull concussive model of a controlled impact on the skull at bregma(*14-16,32*). Mice subjected to this procedure exhibit sparse demyelination in the corpus callosum under the impact site as well as node of Ranvier disruption, which includes exposure and redistribution of axonal K_V_ channels at 7 dpi(*15,16*). Acute treatment with low-dose 4AP starting 24h after concussive TBI was shown to decrease axon damage and demyelination in this mouse model(*16*), supporting the potential use of [^18^F]3F4AP as a companion diagnostic. Compared to sham controls, closed skull TBI mice exhibited higher tracer uptake in the frontal and superior parts of the brain, displaying a diffuse pattern (**Fig. 4A**). To quantify the tracer uptake in this model, an ROI was drawn in the center of the corpus callosum using an atlas (**Fig. 4B**), and the cerebellum ROI was used for normalization across animals Quantification of the PET signal at the injury relative to the cerebellum from 0 to 10 min post injection showed no changes between TBI and sham animals (n = 4-5, per group), indicating no changes in blood perfusion in this area (**Fig. 4C**, left plot). However, signal quantification of from 10-30 min post injection-revealed a higher [^18^F]3F4AP uptake in the TBI animals (**Fig. 4C**, right plot), demonstrating increased [^18^F]3F4AP binding in this location likely due to node of Ranvier disruption and/or demyelination. LFB staining of brain tissue sections from these mice showed that [^18^F]3F4AP was able to detect subtle white matter pathology in the absence of overt tissue cavitation or evidence of focal lesions in the target region (**Fig. 4D**).

**Figure 4.**
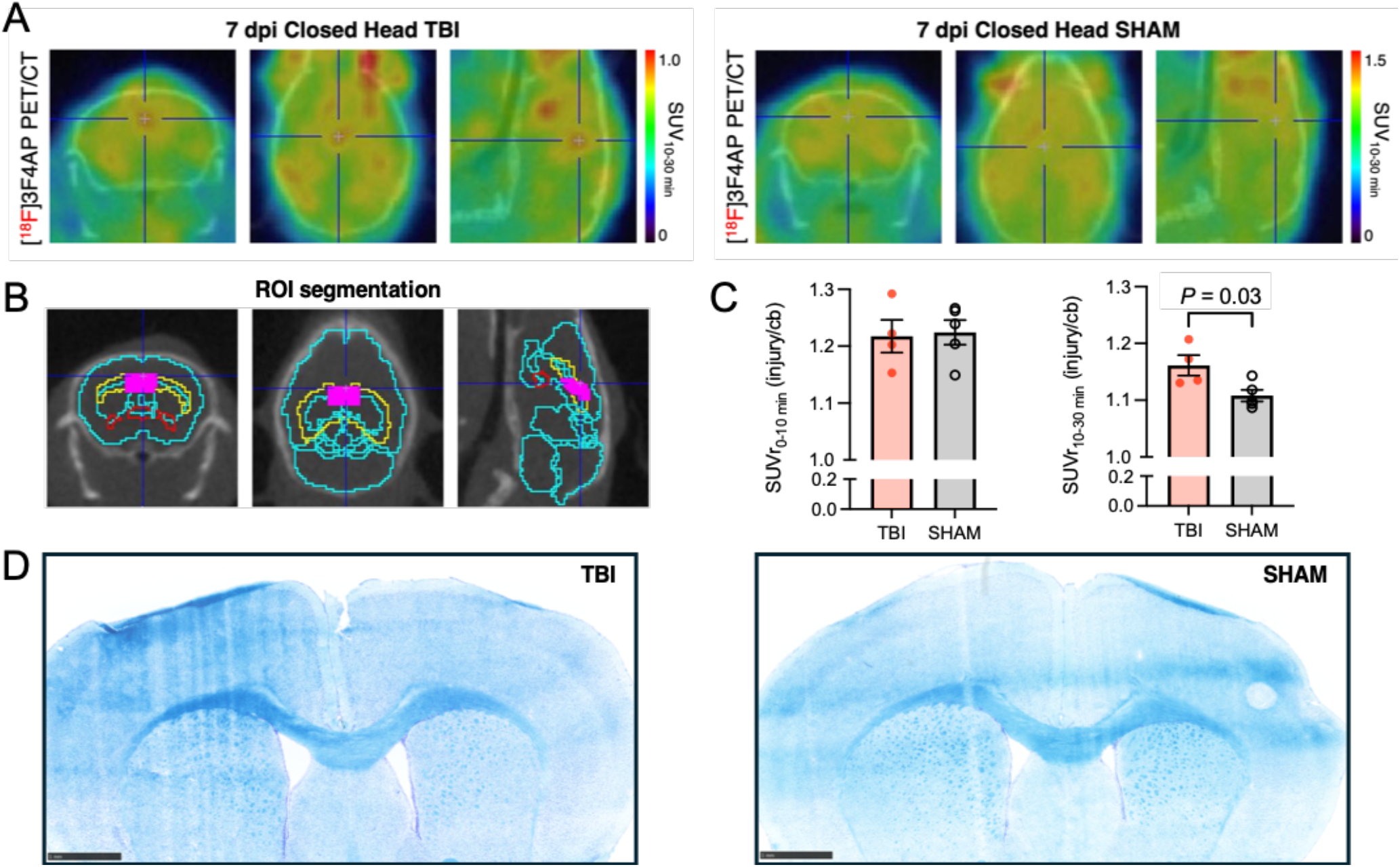
Evaluation of [^18^F]3F4AP in a mouse model of non-penetrating TBI. **A**. Representative coronal, horizontal and sagittal views of brain [^18^F]3F4AP PET/CT in a closed head TBI and sham mice at 7 dpi. Crosshairs are positioned on injury ROI (corpus callosum (CC) directly under bregma). **B**. MRI atlas-guided segmentation for ROI segmentation: injury ROI (pink) was positioned in the coronal slice containing both the CC (yellow) and anterior commissure (red). **C**. Quantification of standardized uptake value in early (SUVr_0-10 min_(injury/cb)) and late PET frames (SUVr_10-30 min_(injury/cb)) for TBI (*n* = 4) and sham (*n* = 5) mice at 7 dpi. Statistical analyses were performed using an unpaired two-tailed t-test. **D** Selected LFB staining of a TBI and a sham mouse brain at 7 dpi. Grouped data are mean ± SEM.

## DISCUSSION

Clinical assessment of axon and myelin pathology in patients with TBI is notoriously difficult. The lack of sensitive imaging biomarkers to differentiate TBI pathophysiology makes it challenging to diagnose and monitor progression or recovery after TBI. Our study investigates the potential use of [^18^F]3F4AP, a novel PET tracer that targets exposed K^+^ channels in demyelinated axons, as a valuable biomarker to expand *in vivo* assessments for TBI. Imaging axons with node of Ranvier disruption and demyelination after TBI is important for detecting axonal injury that is potentially reversible(*11*). This early stage of axon damage is most likely to respond to interventions and warrants more extensive efforts to develop treatments that promote remyelination to protect axons and recover function.

A previous study in a monkey with an accidental open-head TBI incurred during a surgical procedure showed a large increase in [^18^F]3F4AP uptake in the injury site even 3 years after the injury. This injury was not detectable with other common tracers such as [^18^F]FDG (metabolism), [^11^C]PBR28 (TSPO/inflammation) or [^11^C]PIB (amyloid), underscoring the unique potential of [^18^F]3F4AP for imaging chronic TBIs. Since this type of injury could not easily be replicated in monkeys, we set out to study an open head penetrating injury model in mice. [^18^F]3F4AP PET after a CCI penetrating injury showed the highest focal uptake in the impacted area 7 days post injury, compared to the contralateral control area, as well as other brain regions. This increased PET uptake was not observed in sham mice. Longitudinal imaging in these mice showed a moderate increase at 3 dpi, suggesting partial myelin detachment and early changes in K^+^ channel distribution. The signal peaked at 7 days post injury, consistent with peak of demyelination, remained elevated at 14 dpi and gradually by 31 dpi, likely indicating myelin repair and/or axonal loss. LFB staining for myelin provided qualitative confirmation of these changes. The observed [^18^F]3F4AP increase and time course closely aligns with a prior PET, autoradiography and immunohistochemistry (IHC) study in rats after spinal cord injury, which conclusively showed that [^18^F]3F4AP was binding to demyelinated axons(*28*). Furthermore, these findings are also in agreement with the prior results in monkey with TBI, supporting the validity of the mouse CCI model for studying TBIs in primates.

Following the positive findings in the penetrating injury model, we examined a model of non-penetrating TBI, which more closely resembles the predominant TBI type in humans(*33*). In this closed-skull injury model, [^18^F]3F4AP showed a smaller and more diffuse increase in uptake, consistent with the published characterization of a milder and more widespread pathology(*14-16*). Recent findings in this model show that TBI induces node of Ranvier disruption and demyelination, which are reduced by 4-AP treatment during the first week post-injury(15). In this context, our findings suggest that [^18^F]3F4AP could be used to assess axon damage and demyelination after TBI and potentially inform treatment response. Furthermore, there is a pressing need for biomarkers that can detect axon and myelin integrity in chronic phase TBI, adding to the potential impact of [^18^F]3F4AP.

Despite the promising findings, several limitations must be considered. Only female mice were used in the penetrating model and only male mice were used in the non-penetrating model limiting the study of sex as a biological variable. In the closed-head injuries, we only examined 7 dpi, raising the question of whether [^18^F]3F4AP can detect changes over time to monitor the trajectory of progressive degeneration after TBI. We used SUV from 10-30 min post injection as a proxy for tracer binding due to the difficulty of doing kinetic modeling in mice. While this approach appears robust and is justified based on prior pharmacokinetic modeling in monkeys, it is not a direct measure of binding. Furthermore, no clinical *in vivo* scores were available for correlation, which would have strengthened the translational impact of the study. Lastly, PET resolution in the mouse brain is inherently constrained, which made it difficult to clearly visualize the inconspicuous lesion in the closed head injury.

Overall, our findings highlight the utility of [^18^F]3F4AP PET in detecting and monitoring pathophysiological effects of TBI. The tracer exhibited large increases in binding after focal penetrating injuries, likely due to greater levels of demyelination, while non-penetrating injuries produced a weaker but still detectable increase. Future work should focus on evaluating this tracer in human TBI given the dire need of quantitative biomarkers in TBI that can improve diagnosis and be used to monitor new treatments.

## ACKNOWLEDGEMENTS

We thank David Lee and Kyle Stewart at the MGH PET cyclotron facility for producing fluorine-18, Tricia Lacefield and the MGH CCM animal facility staff for rodent transfer and handling.

## AUTHOR CONTRIBUTIONS

Conceptualization: KMRT, PB

Methodology: KMRT, RCA, PB

Investigation: KMRT, KZ, KLR, ETM, LZ, YPZ, EHL, RCA

Visualization: KMRT, LZ, PB

Formal Analysis: KMRT, LZ, KLR, PB

Data Curation: KMRT, RCA, PB

Resources: KMRT, EHL, RCA, PB.

Validation: KMRT

Funding acquisition: KMRT, PB

Project administration: KMRT,

PB Supervision: EHL, RCA, PB

Writing – original draft: KMRT, PB

Writing – review & editing: ALL

## DISCLAIMER

PB is a named inventor on patents related to [^18^F]3F4AP owned by the University of Chicago. Dr. Brugarolas’ interests were reviewed and are managed by MGH and Mass General Brigham in accordance with their conflict-of-interest policies. The other authors declare no conflict of interests.

## DATA AVAILABILITY

The datasets generated and/or analysed during the current study are available from the corresponding author upon reasonable request.

